# Social transmission of egg rejection in a cuckoo host

**DOI:** 10.1101/2020.11.16.384503

**Authors:** Canchao Yang, William E Feeney

## Abstract

Social learning can enable the rapid dissemination of behaviors throughout a population. Rejection of foreign eggs is a key defense in hosts of avian brood parasites; however, whether social cues can inform whether a host rejects an egg remains unknown. Here, we aimed to determine whether access to social information can influence egg rejection behavior in semi-colonial barn swallows (*Hirundo rustica*). By manipulating the social information available from a neighboring nest, we found that swallows that had access to social information (i.e. neighbor recently rejected an egg) were more likely to reject a foreign egg compared to those that did not have access to social information (i.e. neighbor did not reject an egg). This result provides the first empirical evidence that egg rejection behavior can solely be informed by social information, and in doing so highlights the dynamic nature of defenses that hosts can deploy against brood parasitism.

## Introduction

Species interactions are important ecological processes that shape and regulate biodiversity (Thompson 1999). Phenotypes that underpin an individual’s ability to survive and reproduce can emerge and spread throughout a population via several processes, including those that refine over generations (e.g. morphological characteristics) and are passed on through genetic inheritance (Phillips & Shine 2004; Spottiswoode & Stevens 2012), and those that emerge rapidly (e.g. behavioral innovations) and can be spread throughout a population through social transmission (Thornton & Raihani 2008; Davies & Welbergen 2009; Aplin *et al*. 2015). The tractable “arms-races” that emerge between avian brood parasites and hosts has contributed to their use as model systems to study the ecology and evolution of species interactions (Feeney *et al*. 2014; Soler 2014). Unlike most birds, brood parasites lay their eggs in the nests of other birds and foist the burden of parental care onto the host. The cost of brood parasitism selects reciprocal defensive adaptations in hosts and offensive adaptations in parasites that span all stages of the host’s nesting cycle (Brooke & Davies 1988; Langmore *et al*. 2003; De Marsico *et al*. 2012; Feeney *et al*. 2012).

Of the various defenses that hosts deploy against brood parasitism, rejection of foreign eggs is best studied (Soler 2014). The decision to reject an egg is enabled through well-defined behavioral “rules”, such as rejecting the odd egg out (the ‘discordancy hypothesis’) or rejecting any egg (or eggs) that do not match a learned or innate internal template (‘true recognition’) (Rothstein 1974; Lotem *et al*. 1995; Hauber & Sherman 2001; Stevens *et al*. 2013). The decision to reject an egg can be further influenced by personal information about a host’s perceived risk of parasitism (Davies *et al*. 1996; Feeney *et al*. 2015), and once egg rejection emerges in a population it can persist despite generations of allopatry with brood parasites (Lahti 2006; Hale & Briskie 2007; Peer *et al*. 2011). However, despite evidence of social transmission of information being important for other defenses against brood parasitism, such as recognition and mobbing of adult parasites (Davies & Welbergen 2009; Feeney & Langmore 2013), and suspicion that the decision to reject an egg may too be impacted by social information (Soler 2011), empirical evidence that egg rejection can be influenced by social information is lacking.

Here, we used the barn swallow (*Hirundo rustica*) to investigate whether egg rejection behaviors are influenced by access to social information. In China, barn swallows are occasionally parasitized by the common cuckoo (*Cuculus canorus*) with a parasitism rate of 0–2.4% (Yang *et al*. 2015; Su *et al*. 2017), and they are capable of recognizing and grasp-ejecting non-mimetic parasite eggs at a rate of 24–63% (Yang *et al*. 2015). They are semi-colonial (i.e. females nest between 0.5 – 10m from their closest neighbor) and build open cup-shaped nests, permitting the possible visual transmission of egg rejection between individuals (Møller 2002). The high rejection rate and low natural parasitism rate currently experienced by barn swallows in China, as well as their open-cup nests and semi-coloniality, makes them a well-suited model organism to investigate whether egg rejection can be influenced solely by social information.

## Results and Discussion

In 2018, we first conducted a series of egg rejection trials using a blue egg (*n*=55, common cuckoos often lay blue eggs in China (Fossøy *et al*. 2016)), conspecific swallow egg (*n*=48) and no egg (*n*=43, during these trials experimenter behavior at the nest was identical but no egg was added) and found that swallows rejected 61.82% of blue eggs, with 85.29% of rejections occurring within 5h. No eggs were rejected (i.e. ejected or deserted) during the conspecific and no egg trials. Then, in 2019, swallow nests were experimentally set up in pairs 3m apart with nest entrances facing one another (Fig. 1A). Once colonized, a blue egg was placed in one nest (partner nest) from each pair and the behavioral response after 5h was used to determine whether the nest it was paired with (focal nest) would subsequently be used for a “transmission” trial (if the egg was rejected) or “non-transmission” trial (if the egg was not rejected). Nest pairs designated for control trials experienced comparable disturbance to the transmission and non-transmission nests, but no egg was added. Following this, the focal nest received a blue egg during early incubation to investigate whether access to social information increased the likelihood of the swallow from the focal nest rejecting the foreign egg.

**Figure 1.**
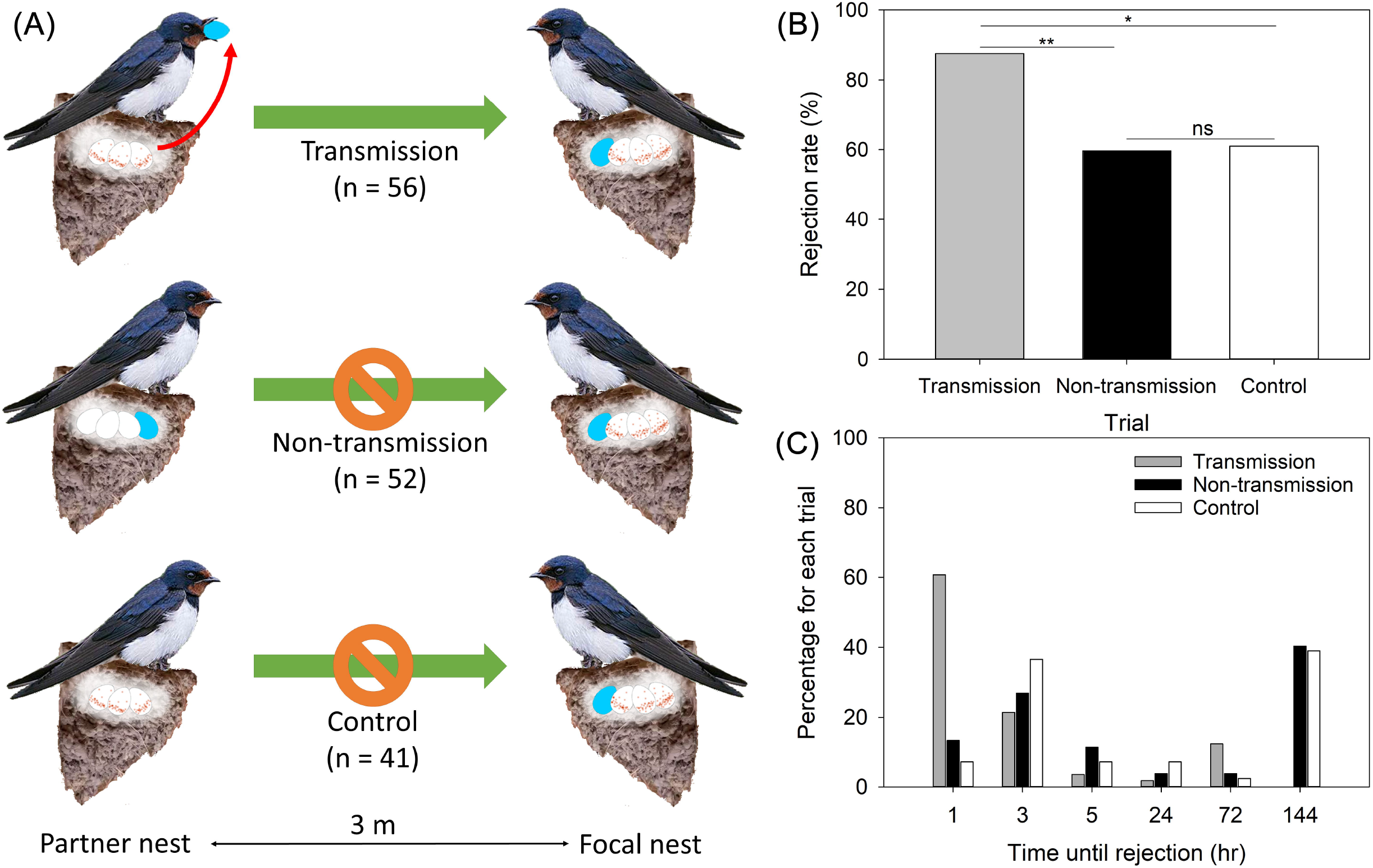
Schematic outlining the experimental design (A): in the transmission trial the focal received a model egg in its nest after the partner nest had rejected a model egg (*n*=52); in the non-transmission trial the focal nest received a model egg after the partner nest had accepted a model egg (*n*=56); and in the control trial the focal nest received a model egg but the focal nest had not previously received a model egg (*n* 41). Rejection and acceptance rates in the transmission, non-transmission and control trials (B). Egg rejection rates across time periods by birds in focal nests across the transmission, non-transmission and control trials (C).

Our results indicate that barn swallows that have access to social information are more likely to reject a foreign egg, and do so more quickly than those that have not had access to social information. We found that there was a significant effect of treatment on egg rejection rates (GLMM: F_2, 137_=4.855, *P*=0.009; Fig. 1B): rejection of blue eggs was significantly higher in the transmission (87.5%, *n*=56) versus non-transmission (59.62%, *N* = 52) (Sequential Bonferroni: *P* = 0.004) trials, and we found no difference in rejection rates between the non-transmission and the control trials (60.98%, *n*=41) (Sequential Bonferroni: *P*=0.839). We also found no significant effects of clutch size (GLMM: F_4,137_=0.767, *P*=0.549) or incubation day (i.e. the number of days after incubation commenced that the trial began) (GLMM: F5,137=0.409, *P*=0.842). We found that time until rejection was significantly predicted by treatment (Cox regression: Wald=10.847, df=2, *P*=0.004), with blue eggs more likely to be rejected within 1h during the transmission trial (60.71% of rejected eggs) compared to the non-transmission and control trials, which were more likely to rejected within 1-3h (13.46% and 7.32%, respectively) (Fig. 1C). Again, we found no effect of clutch size (Cox regression: Wald=1.980, df=4, *P* = 0.739) or incubation day (Cox regression: Wald=2.986, df=5, *P*=0.702) on the time until rejection across treatments.

These data indicate that hosts can mediate egg rejection behaviors solely based off access to social information. It is well-established that defenses that operate to prevent parasitism from occurring, such as recognizing and responding to the threat post by an adult parasite near a nest, are learned behaviors (Davies & Welbergen 2009; Langmore *et al*. 2012; Feeney & Langmore 2013) that wane in the absence of parasitism (Briskie *et al*. 1992; Hobson & Villard 1998; Lindholm & Thomas 2000; Hale & Briskie 2007). Likewise, personal information about the perceived risk of parasitism can affect the likelihood of hosts rejecting a foreign egg (Davies *et al*. 1996; Feeney *et al*. 2015; Thorogood & Davies 2016) or chick (Langmore *et al*. 2009). During the two years over which this study was conducted no natural parasitism was recorded, only blue model eggs were rejected, and the only significant predictor of increased egg rejection was having a neighbor that previously rejected an egg. This suggests that in addition to being highly sensitive to information about the presence of cuckoos in the area, hosts can also be sensitive to parasite-specific behaviors in conspecifics, to the extent that they can make egg rejection decisions solely using this information. The nuanced use of both personal and social information by hosts has implications for coevolutionary trajectories in these systems, potentially underpinning a dynamic mosaic of phenotypically plastic defenses across host populations that parasites must counter, which may outpace and shape genetic evolution.

## Materials and Methods

### Study area and species

In this study, we used a barn swallow (*Hirundo rustica*) population in the Jinping District of southeast China (23° 352 ′-24° 401′ N, 116° 658′-116° 706′ E) to examine whether egg rejection behaviors could be influenced by social information. Barn swallows are insectivorous, semi-colonial passerines that build mud nests on walls under eaves and typically nest within 0.5–10m of their nearest conspecific neighbour. They are occasionally parasitized by the common cuckoo (*Cuculus canorus*), with a parasitism rate of 0–2.4%. Barn swallows are capable of recognizing and grasp-ejecting non-mimetic parasite eggs, with a rejection rate of 24–63% (Yang *et al*. 2015). All experiments in this study were conducted between March–May of 2018 and 2019.

### Experimental procedure

In 2018, we tested the egg rejection rate of the barn swallow population for non-mimetic and conspecific eggs using artificial parasitism experiments. For the non-mimetic (“parasite”) trials, we used blue model eggs that were made of polymer clay and similar in size to barn swallow eggs. The model eggs were blue because while the egg phenotype of common cuckoos that parasitise barn swallows in China is not known, common cuckoos often lay blue eggs in China (Fossøy *et al*. 2016). In each trial, one blue model egg was inserted into each swallow nest on the day following clutch completion. The artificially parasitized nests were checked at 1hr, 3hr, 5hr, 24hr, 72hr, and 144hr after egg placement (6 days in total). In the conspecific trial, the experimental procedure was identical, except that a conspecific egg from another nest was added to the host nest instead of a model egg. All conspecific eggs were returned to the appropriate nest after the experiment. A control treatment, which followed identical experimental procedures except that no egg was added, was performed to control for the effects of disturbance and egg manipulation. We found that the egg rejection rate for this population of the blue eggs was 61.82%, with 85.29% of the rejections occurring within 5hr (*n*=55). All conspecific eggs were accepted by hosts (*n*=48), and no desertions or ejections were observed in any trials, including the control trial (*n*=43). The period from spanning the beginning of egg incubation to the end of nestling feeding was approximately 38 days.

In 2019, we experimentally manipulated the social transmission of egg rejection using the same barn swallow population. Swallow nests were collected before the breeding season (late February) and glued to walls in pairs. The paired nests were 3m apart, with the nest entrances facing each other (Fig. 1A). Each pair of nests was at least 30m from the next closest pair. To ensure that nests were not resampled during the experiment, we required that the nesting period of all pairs included in the study were conducted within a 38-day period. Within each pair of nests, one was considered the focal nest, and the other nest was considered the partner nest.

During the experimental trials, one blue model egg (identical to those used in 2018) was inserted into the partner nest during the early incubation period (i.e. within six days of clutch completion) between 6:30–7:30am. To control for the effects of nest manipulation, the partner nests in the control trials were manipulated, but no eggs were added. To avoid host observation of artificial parasitism, a black umbrella was used as a shield during egg transfers and nest manipulation. After 5hr, the experimental nests were checked; we chose to check the nests after 5hr because most (85.29%) of the rejections in 2018 occurred within 5 hr.

If the egg was not rejected from the partner nest within 5hr, the model egg was removed and the pair of nests was assigned to the non-transmission trial. If the egg was rejected from the partner nest within 5hr, we assigned the pair of nests to the transmission trial. Following this, a blue model egg was inserted into the focal nests of all pairs assigned to the non-transmission, transmission, and control trials (Fig. 1). The focal nests in all three trials were checked 1hr, 3hr, 5hr, 24hr, 72hr, and 144hr after artificial parasitism. The control trial was used to control for the effects of the partner nest on rejection behavior. The focal host response to the model egg was classified as rejection if the parasite egg was ejected or deserted, and as acceptance if the parasite egg continued to be incubated. We predicted that the egg rejection rate in the focal nest would be higher in transmission trial because this individual had previously had access to social information of egg rejection from the partner nest. We also predicted that the egg rejection rats of the focal nests would be similar between the non-transmission and control treatments, as these individuals had not previously had access to social information.

### Statistical analyses

A generalized linear mixed model (GLMM) was used to quantify the effects of artificial parasitism on egg rejection among trials. In this model, the rejection or acceptance of the parasite egg was set as the response variable; the three trials (non-transmission, transmission, and control), clutch size, and incubation day (i.e. the number of days after incubation commenced that the trial began) were set as fixed effects; and nest identity and date of first egg laying date were set as random effects. We compared pairs of significant factors using Sequential Bonferroni tests. Cox regressions, which calculate survival probabilities at given time points, were used to compare the speed of egg rejection among trials. That is, we evaluated the probability that the parasite egg had not yet been rejected at each time point (1hr, 3hr, 5hr, 24hr, 72hr, and 144hr) for each trial. Thus, this analysis compared how quickly the hosts in different trials recognized the parasite egg. All statistical analyses were performed using IBM SPSS 25.0 for Windows (IBM Inc., USA).

## ACKNOWLEDGEMENTS

This work was funded by the Natural Science Foundation of China (No. 31672303) (CY) and the Alexander von Humboldt Foundation (WEF).

## AUTHOR CONTRIBUTIONS

CY conceived the idea, with assistance from WEF. CY collected and analyzed the data. Both authors wrote the manuscript.

## DATA AVAILABILITY

All data used in this manuscript is provided in the supplementary materials (Supplemental File 1).

